# Nuclear Receptor 5A2 Regulation of *Agrp* underlies Olanzapine-induced Hyperphagia

**DOI:** 10.1101/2022.02.15.480590

**Authors:** Rizaldy C. Zapata, Dinghong Zhang, Avraham Libster, Alessandra Porcu, Patricia Montilla-Perez, Zhi Zhang, Stephanie M Correa, Francesca Telese, Olivia Osborn

## Abstract

Antipsychotic (AP) drugs are highly efficacious treatments for psychiatric disorders, but a serious side effect of their use is excessive weight gain and subsequent development of metabolic disease. Increased food intake is the underlying driver of AP-induced weight gain, although the underlying mechanisms remain unknown. In previous studies, we identified hypothalamic genes whose expression level was altered following APs-induced hyperphagia. Among these genes, the orexigenic peptide Agrp and the transcription factor nuclear receptor subfamily 5 group A member 2 (Nr5a2) were two of the most significantly upregulated genes by APs. NR5a2 is broadly expressed throughout the body, but little is known about its role in the brain. In this study, we investigated the role of hypothalamic NR5a2 in AP-induced hyperphagia and weight gain. In hypothalamic cell lines, OLZ treatment resulted in a dose-dependent increase in gene expression of *NR5a2* and *Agrp*. In mice, administration of a specific Nr5a2 inhibitor decreased olanzapine-induced hyperphagia and weight gain, while knockdown of Nr5a2 in the arcuate nucleus (ARC) partially reversed olanzapine-induced hyperphagia. Chromatin-immunoprecipitation-PCR studies showed for the first time that NR5a2 directly binds to the *Agrp* promoter region. In addition, *in situ* hybridization studies confirm that *NR5a2* and *Agrp* are co-localized in a subset of cells in the arcuate nucleus. In summary, we identify Nr5a2 as a key mechanistic driver of AP-induced food intake and these findings can be used to inform future clinical development of APs that do not activate hyperphagia and weight gain.

## INTRODUCTION

Antipsychotic (AP) medications are highly efficacious treatments for various psychiatric disorders^1–6^ but a serious side effect of their use is excessive weight gain^3,7,8^. Approximately 20% of patients treated with a broad range of APs gain clinically significant amounts of weight (>7% of their baseline weight)^9^. Drug safety reviews have shown the percentage of patients gaining a clinically significant amount of weight varies between individuals and depending on the drug, ranging from ~20-40% for olanzapine (OLZ) and clozapine and ~10-20% for quetiapine and risperidone^9–16^. While OLZ is associated with a very high risk for weight gain, it is also regarded as one of the most clinically effective medications^17^. APs induce weight gain in both human^7,18–20^ and rodents ^21–26 27–30^, by increasing food intake (hyperphagia). Despite widespread efforts to understand how APs induce hyperphagia, very little is known about the mechanisms underlying this serious adverse effect. Previous studies have relied on non-specific anti-obesity drugs that suppress basal feeding to reduce AP-induced weight gain (i.e. locaserin^31^, orlistat^32^, liraglutide^33^, nizatidine^34^ metformin^35^). While using anti-obesity drugs in combination with APs is clinically beneficial to offset weight gain, they do not shed light on the specific mechanisms underlying AP-induced hyperphagia. Delineating the specific mechanisms driving AP-induced hyperphagia can be used to inform future drug development of highly effective APs without this metabolic liability and also more broadly to understand pathways regulating food intake that could be used as potential anti-obesity strategies.

In our previous work, we used a *C. elegans*-based high-throughput screen^36^ to identify specific chemical suppressors of AP-induced hyperphagia^27^. We then conducted studies in a well-established mouse model of AP-induced hyperphagia and weight gain^26–31,37–40^ to determine whether the compounds identified in the *C. elegans* screen could also suppress AP-induced hyperphagia in mammals^27^. These studies identified hypothalamic genes whose transcriptional levels were associated with APs-induced hyperphagia, which include the orexigenic peptide Agrp and the transcription factor nuclear receptor subfamily 5 group A member 2 (Nr5a2), among the most significantly upregulated genes by APs. Food intake is regulated by many parts of the brain^41^ and agouti-related peptide expressing (*Agrp*) neurons in the arcuate nucleus (ARC) of the hypothalamus play a major role^42–46^. While some studies have reported increased expression of *Agrp* after AP-treatment^26,47,48^, the importance of this neuronal subtype and the molecular mechanisms regulating the AP-induced expression of this key pro-feeding gene are not well understood^49–51^. Furthermore, the hypothalamic upregulation of Nr5a2 is specifically associated with AP treatment and not increased body weight, suggestive of its important role in AP-induced hyperphagia^27^.

*Nr5a2*, also called Liver receptor homolog −1 (LRH-1), is best known for its role in the periphery and governs a transcriptional network of genes involved in bile acid signaling and liver lipid homeostasis^52–54^. Nr5a2 has also been implicated in adipocyte formation^55^, intestinal function^56^ pancreatic inflammation^57^ and expression of pancreatic digestive enzymes^58,59^. However, Nr5a2 is also widely expressed in the mouse brain^60^ where it controls neural stem cell fate^61^. Within the brain, Nr5a2 is expressed in the ARC of the hypothalamus^62–64^ and more recently has been shown to mark a select subset of neurons in this region^65^ but little is known about its role in the central nervous system (CNS)^61,64^. Our previous studies provided the first insights into the potential involvement of NR5a2 in AP-induced food intake. In these *C.elegans* based studies, we determined that NR5a2 ortholog/nhr-25 mutant strain (nhr-25(ku215)) was resistant to AP-induced hyperphagia^27^. NR5a2 is broadly expressed throughout the body and has well-described roles in the liver ^54,66^, gut ^56^, and pancreas ^57,67^ but little is known about its role in the CNS^61,64^. In the current study, we used several mouse models to investigate the role of NR5a2 in antipsychotic-induced food intake and weight gain.

## MATERIALS AND METHODS

### In vitro studies

Adult mouse hypothalamic cell lines (mHypoA-59, CLU468 cells, Cedarlane) were cultured as described previously ^68,69^. In brief, cells were grown and maintained in high-glucose, pyruvate-free DMEM supplemented with 10% fetal bovine serum, L-glutamine (Cat. 25030081, Gibco, NY), and 10 u/ml of penicillin and 10 ug/ml of streptomycin (Cat. 15149-122, Gibco) of in a 5% CO_2_ environment. Cells were treated with OLZ (25μM–200μM) for 6 hours and then RNA extracted for gene expression analysis. Cells were also co-treated with NR5a2 antagonist SR1848 (Aobious, Gloucester, MA) for 6 hours at 1-5μM ^70^ in DMSO as described previously.

### Gene expression

RNA isolation was performed using Trizol (cat # 15596026, Invitorgen) and was purified using RNeasy Plus Mini Kit (cat # 774104, Qiagen) using the manufacturer’s recommendations. cDNA was reverse transcribed from 300 ng of RNA using High Capacity cDNA transcription kit (cat # 4368813, Applied Biosystems). Relative expression analyzed by qPCR using StepOne Realtime PCR System. Gene expression was calculated after normalization to the housekeeping genes^71^ using the Δ^ΔCt^ method. Gene expression was calculated relative to experimental controls. Primer sequences (5’-3’) used to measure gene expression are listed in Table 1.

**Table 1.**
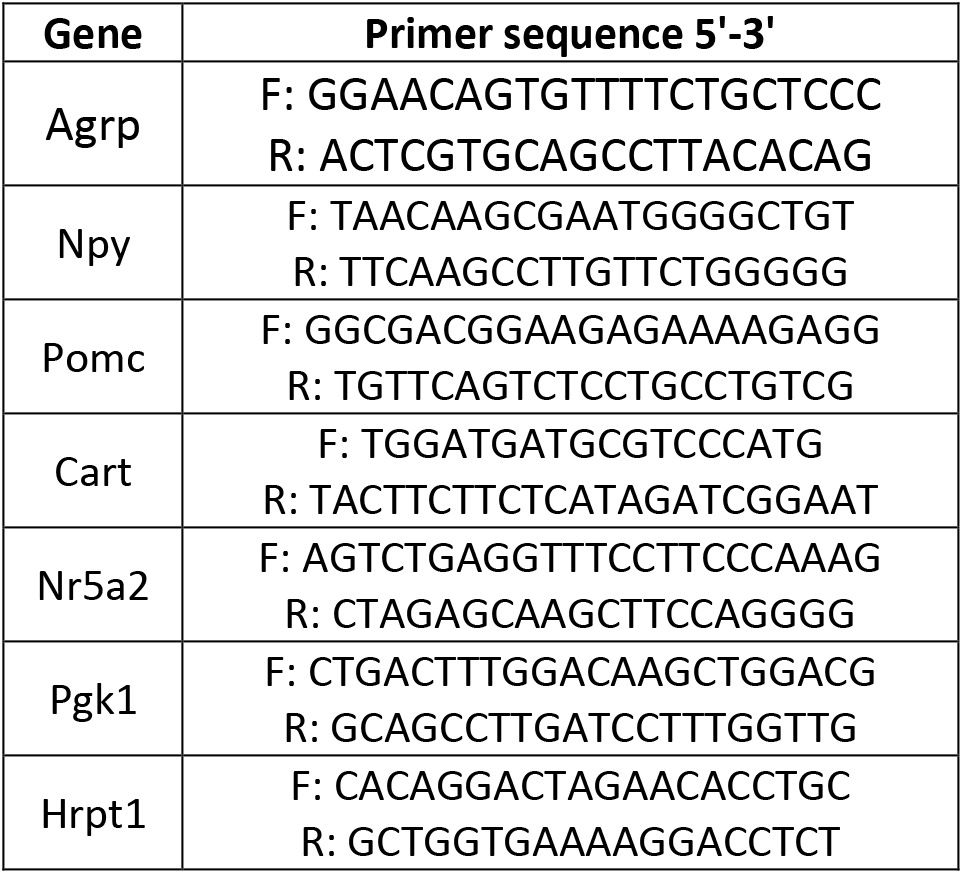

### Mice

All protocols were approved by UCSD IACUC. All mice were singly housed in standard cages and acclimated to laboratory conditions (12:12 light-dark, 20-21°C, 50% humidity) for 7 days before experimentations. Mice were singly housed to allow accurate measurement of daily food intake by weighing food in the hopper and accounting for any spillage^72^. All studies were performed in female C57B6/J mice (Jackson, stock # 000664) or *Agrp* null mice. *Agrp*^−/−^ mice were gifted by Dr. Chen Liu of UT Southwestern.

### Olanzapine administration

Olanzapine (OLZ) was compounded into 45% HFD diet (54 mg/kg =~6-8mg/kg) as a convenient dosing strategy^27,29,30^, and this approach has been used in many other studies investigating AP-induced hyperphagia and weight gain^26,37–40^. This dose results in mouse plasma levels (21±5 ng/ml) that are similar to levels observed in humans treated with OLZ (10–50 ng/mL)^38^.

### Systemic inhibition of Nr5a2

Twelve-week old female mice were acclimated to receive intraperitoneal (IP) injections of sterile saline for 3 days before the experiment and then were randomized to receive a 45% high fat diet (CON, D09092903B, Research Diets) with or without olanzapine (OLZ, 54mg/kg, D16111030). Mice were then further randomized to receive the vehicle solution (VEH, 10% DMSO, 10% Tween 80 in 0.9% NaCl) or the Nr5a2 inhibitor (SR1848) at 30 mg/kg daily for 7 days. Food intake and body weight were measured daily. Animals were sacrificed at the end of the study and the hypothalamus was dissected, snap frozen in liquid nitrogen, and stored in −80°C until analyses.

### Hypothalamic inhibition of Nr5a2

Twelve-week old female mice were anesthetized with isoflurane and were mounted on a heating pad on a Neurostar robotic stereotaxic surgery set-up. Nr5a2 siRNA (SMARTpool: Accell, Catalog ID:L-047044-01-0005, Dharmacon, Lafayette,CO) or non-targeting control siRNA (Catalog # K-nin; Dharmacon, Lafayette,CO) (n=4-5) was delivered bilaterally into the ARC using coordinates: A-P: −1.58 mm from Bregma; M-L ± 0.25 mm from midline; D-V: −5.8 mm into the skull. Mice were allowed to recover for 7 days before transitioning to CON or OLZ. Food intake was measured daily and body weight every other day for 14 days. Animals were sacrificed at the end of the study and the hypothalamus was dissected, snap frozen in liquid nitrogen, and stored in −80°C until analyses.

### *Agrp* null studies

Twelve-week old WT and *Agrp*^−/−^ female mice were randomized to receive either CON or OLZ (n=9-17/group). Food intake was measured daily and body weight every other day for 12 days. Animals were sacrificed at the end of the study and the hypothalamus was dissected, snap frozen in liquid nitrogen, and stored in −80°C until analyses.

### Chromatin Immunoprecipitation (ChIP)

ChIP experiments were conducted in triplicates using methods previously described in other neuronal cell types ^73–75^. Briefly, hypothalamic mHypoA-59 cells were grown in 10cm dishes and at 75-80% confluency and fixed with 1% formaldehyde. Nuclei were isolated before chromatin extraction. Chromatin from approximately 10 million cells was sheared using a sonication device (Bioruptor Pico, #2013-2019, Diagenode) and optimized to produce ~400bp fragments. Chromatin was immunoprecipitated using 4ug of Nr5a2 antibody (PP-H2325-00. 5 μg/ChIP, RD Biosystems)^57^ and 20ul of beads without any antibody were used as control sample. Importantly, this antibody has been validated in the Nr5a2 KO^52^ and has successfully been used in liver^76,77^ and pancreatic^57^ ChIP experiments in mice. After primary and secondary antibody incubation and washes, purified DNA was used in quantitative PCR reactions with primers targeting the promoter of NR5a2 target gene Prospero Homeobox 1(Prox1) promoter (Prox1-F 5’-CTGTTAACTGTGCCCAGGGAGAGGA-3’, Prox1-R 5’-TGGTTTGACATCTTGGGTGA-3’) ^61^ as a positive control^61^ or the *Agrp* promoter region (*Agrp*-F 5’-GGGGTCTGGACACCCTATCT-3’, *Agrp*-R 5’-CACACGTGACTGCTTCCTGT-3’)^78^. Fold enrichment was calculated relative to the no-antibody control samples.

### RNAscope

WT female C57BL6 mice were anesthetized with Pentobarbital then transcardially perfused with 20 ml PBS followed by 40 ml 10% Formalin/PBS (Sigma). Brains were removed and incubated in 15% sucrose/10% Formalin overnight at 4°C. Following cryoprotection in 30% sucrose/PBS, brains were embedded in OCT on dry-ice and stored at −80°C. Serial 20 μm sections were cut using a cryostat and mounted on glass slides (VWR) and sections stained with RNAscope^®^ Probe - Mm-Nr5a2-O1-C2 (cat number 547841-C2), *Agrp* RNAscope^®^ Probe - Mm-*Agrp* (cat number 400711).

## RESULTS

### Olanzapine treatment results in significant elevation of *NR5a2* and *Agrp* in hypothalamic cell lines

OLZ treatment of hypothalamic cells resulted in a significant dose-dependent upregulation of both *Nr5a2* (**Fig. 1A**) and *Agrp* genes expression (**Fig. 1B**) compared to vehicle treatment. While in general AP-drugs drive hyperphagia and weight gain, there is significant variation in the magnitude of these effects ^79^. In our recent study,^29^ we measured antipsychotic-induced weight gain (AIWG) in mice and stratified into subgroups that were highly prone to weight gain (gained 6.3g body weight) and weight gain resistant (gained 1.3g body weight) ^29^. In addition to the previously noted elevation in the hypothalamic expression of *Agrp*, we also observed a highly significant elevation of *Agrp* (**Fig. 1C**) and *NR5a2* (**Fig. 1D)** expression in the AIWG-prone compared with the AIWG-resistant mice. In addition, AIWG-prone mice were also hyperphagic compared to AIWG-resistant mice and thus the elevated hypothalamic expression of *NR5a2* and *Agrp* in the AIWG-prone mice further suggests that these genes may play a role in AP-induced hyperphagia and weight gain.

**Figure 1.**
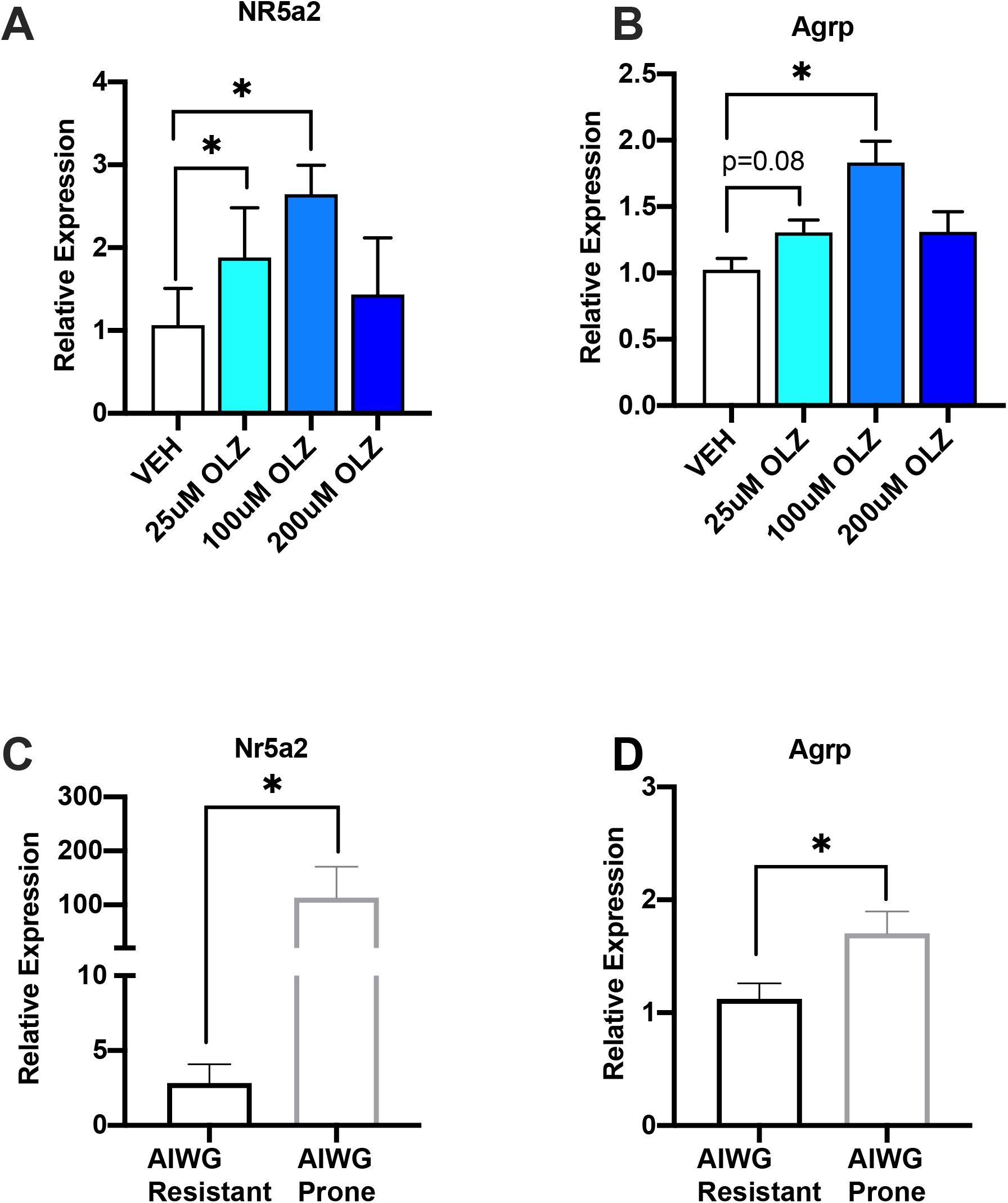
OLZ treatment is associated with elevated *Nr5a2* expression. OLZ treatment of hypothalamic cells results in dose dependent increase in expression of (**A**) *Nr5a2* and (**B**) *Agrp.* Mice that are highly ***Prone*** to Antipsychotic-Induced Weight Gain (AIWG-P) have significantly elevated hypothalamic levels of (**C**) *Nr5a2* and (**D**) *Agrp* compared with AIWG-***Resistant*** mice (AIWG-R). Data is expressed as mean ± SEM and was analyzed using one-way ANOVA followed by uncorrected Fisher’s LSD test, (A-B, (n= 3-8 replicates per group) or students t-test (C-D, n= 8-10 replicates per group), * denotes statistical significance at *p* < 0.05.

### NR5a2 inhibitor treatment reduces AP-induced food intake and weight gain

We further investigated the role of Nr5a2 in AP-induced hyperphagia in mice using a specific Nr5a2 antagonist (SR1848, IP 30mg/kg daily)^70^. SR1848 inhibits NR5a2 function by triggering translocation of Nr5a2 from the nucleus to the cytoplasm, which ultimately abrogates its ability to transduce transcription of its targets^70^. As expected, OLZ treatment resulted in elevated hypothalamic expression of NR5a2 and co-treatment with SR1848 (OLZ+SR) did not impact NR5a2 expression levels (**Fig. 2A**). However, co-treatment of OLZ with SR1848 resulted in significantly reduced daily food intake (**Fig. 2B)** and weight gain (**Fig. 2C**) compared with OLZ alone over 7 days of treatment. Furthermore, hypothalamic levels of *Agrp* (**Fig. 2D**) were significantly reduced by co-treatment, while other appetite regulating neuropeptides *Npy and Pomc* levels were not significantly changed. To determine whether SR1848 has a direct effect on hypothalamic gene expression, we treated hypothalamic cell lines with SR1848 and measured *Agrp* gene expression (**Fig. 2E**). We observed significant reduction in *Agrp* expression levels after either 1uM or 5uM SR1848 dosing for 6 hours suggesting inhibition of NR5a2 in the hypothalamus has a direct impact on *Agrp* gene expression.

**Figure 2.**
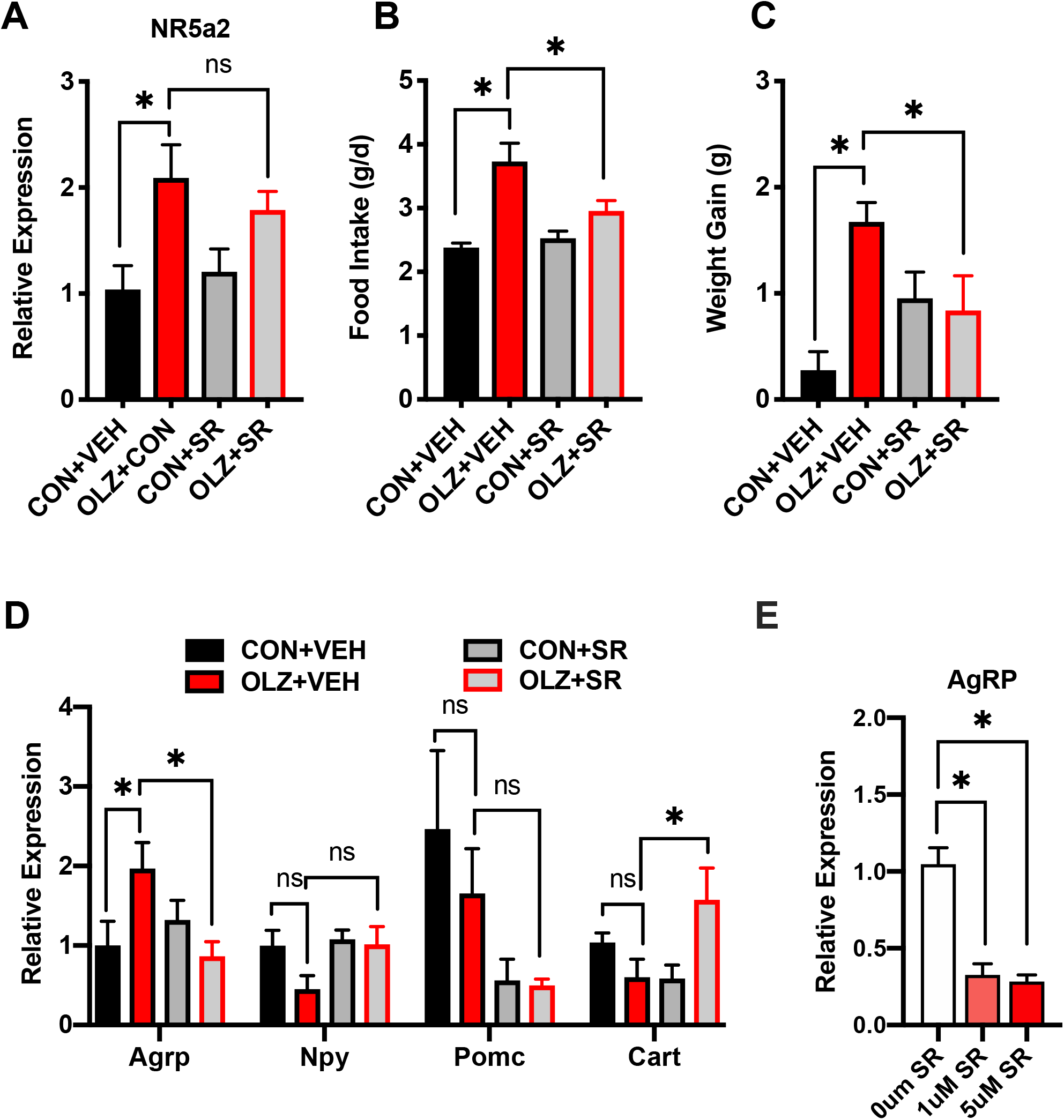
Systemic Nr5a2 antagonist treatment reduces food intake and weigh gain in mice treated with Olanzapine. **A)** *Nr5a2* expression, (**B)** Average daily food intake, (**C**) Weight gain, (**D**) Hypothalamic neuropeptide expression in C57BL/6 WT female mice fed either control diet (CON) or OLZ diet and injected with NR5a2 antagonist (SR1848, ‘SR’, 30mg/kg) or vehicle (VEH) for 7 days. **E**) Quantitative PCR determination of *Agrp* expression in hypothalamic cells lines treated with SR1848 (1uM or 5uM) for 6 hours. Data is expressed as mean ± SEM and was analyzed using one-way ANOVA followed by uncorrected Fisher’s LSD test, * denotes statistical significance at *p* < 0.05, n= 4-11 replicates per group.

### The knockdown of Nr5a2 in the arcuate nucleus partially reversed olanzapine-induced hyperphagia and weight gain

To determine whether AP-induced food intake and body weight regulation *in vivo* require the expression of Nr5a2 specifically in the hypothalamus, we used siRNA-mediated knockdown of Nr5a2 expression in the hypothalamus (**Fig. 3A**). As expected, Nr5a2 expression was increased by OLZ treatment, and treatment with siRNA targeting NR5a2 significantly reduced NR5a2 expression (**Fig. 3A**). OLZ treatment increased food intake (**Fig. 3B**) and body weight gain (**Fig. 3C-D**) which was reversed by hypothalamic NR5a2 siRNA treatment. Furthermore, gonadal (gWAT) and subcutaneous (sWAT) fat mass were also significantly elevated by OLZ treatment and significantly reduced by NR5a2 siRNA compared to siRNA control (**Fig. 3E**).

**Figure 3.**
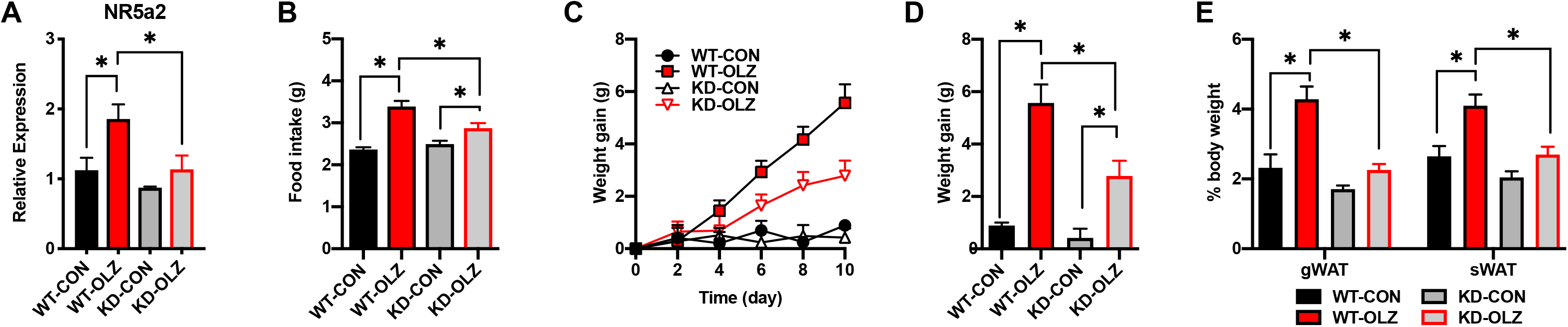
Hypothalamic knockdown of *Nr5a2* significantly bunts OLZ-induced food intake and weight gain. siRNA mediated knock down of *Nr5a2*, delivered by stereotaxic injection to the arcuate nucleus, results in (**A**) reduced expression of *Nr5a2*, (**B**) blunted OLZ-induced food intake and (**C-D**) reduced OLZ-induced weight gain and body fat (**E**), Data is expressed as mean ± SEM and was analyzed using one-way ANOVA followed by uncorrected Fisher’s LSD test, * denotes statistical significance at *p* < 0.05, n=4-5 per group.

### Genetic deletion of *Agrp* in mice prevented olanzapine-induced hyperphagia and weight gain

Since OLZ treatment increases the expression of *Agrp* similarly to Nr5a2, we tested whether *Agrp* is necessary for the hyperphagic effect of OLZ. As expected, OLZ treatment of WT mice induced higher food intake (**Fig. 4A**) and weight gain (**Fig.4B-C**) compared with control-treated mice. On the other end, *Agrp*KO mice were resistant to the hyperphagic and weight gain response to OLZ treatment (**Fig.4A-C**). While OLZ treatment resulted in elevated hypothalamic transcriptional levels of *NR5a2* in *Agrp*KO mice compared with control treated KO mice, the expression of *Npy*, *Pomc* and *Cart* was unchanged, (**Fig 4.D**) suggesting NR5a2 maybe upstream of Agrp regulation.

**Figure 4.**
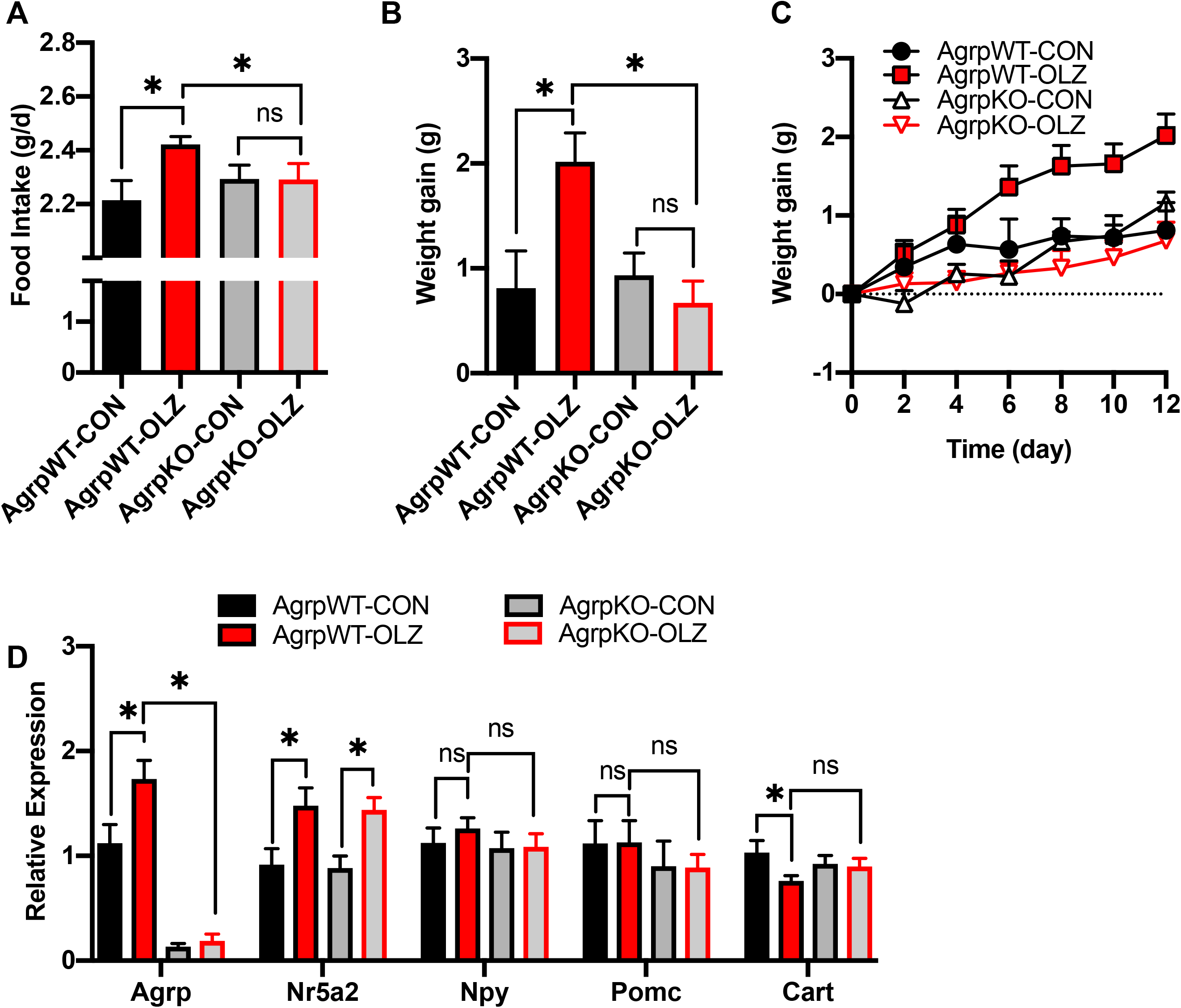
Agrp KO mice are resistant to OLZ-induced hyperphagia and weight gain. **A)** Food intake (**B-C**) weight gain, (**D**) hypothalamic gene expression in WT and KO mice treated with CON or OLZ diets. Data is expressed as mean ± SEM and was analyzed using one-way ANOVA followed by uncorrected Fisher’s LSD test, * denotes statistical significance at *p* < 0.05, n=9-12 per group.

### NR5a2 directly regulated *Agrp* expression

These data led us to hypothesize that NR5a2 may directly regulate the expression of *Agrp* by binding to its promoter. To test this, we conducted chromatin immunoprecipitation with NR5a2 antibodies followed by PCR (ChIP-PCR) in hypothalamic mHypo-59A cells. In agreement with previous studies in neuronal stem cells,^61^ we found that Nr5a2 binds the Prospero Homeobox 1(Prox1) promoter (**Fig. 5A**). We then used primers specific for *Agrp* promoter region ^78^ and determined ~2-fold enrichment of Nr5a2 binding to the *Agrp* promoter region over the control sample (**Fig.5B**). These ChIP experiments in hypothalamic cells identify Agrp as a direct transcriptional target of the transcription factor NR5a2. In situ hybridization analysis determined that NR5a2 is expressed highly and specifically in the ARC of the hypothalamus (**Fig. 5C**). Co-staining with *Agrp* revealed that a subset of NR5a2 expressing neurons also contain *Agrp* (**Fig. 5C**). Therefore, these studies suggest that a refined population of *NR5a2*-expressing cells co-express *Agrp* in the ARC which play a major role in AP-induced hyperphagia and weight gain.

**Figure 5.**
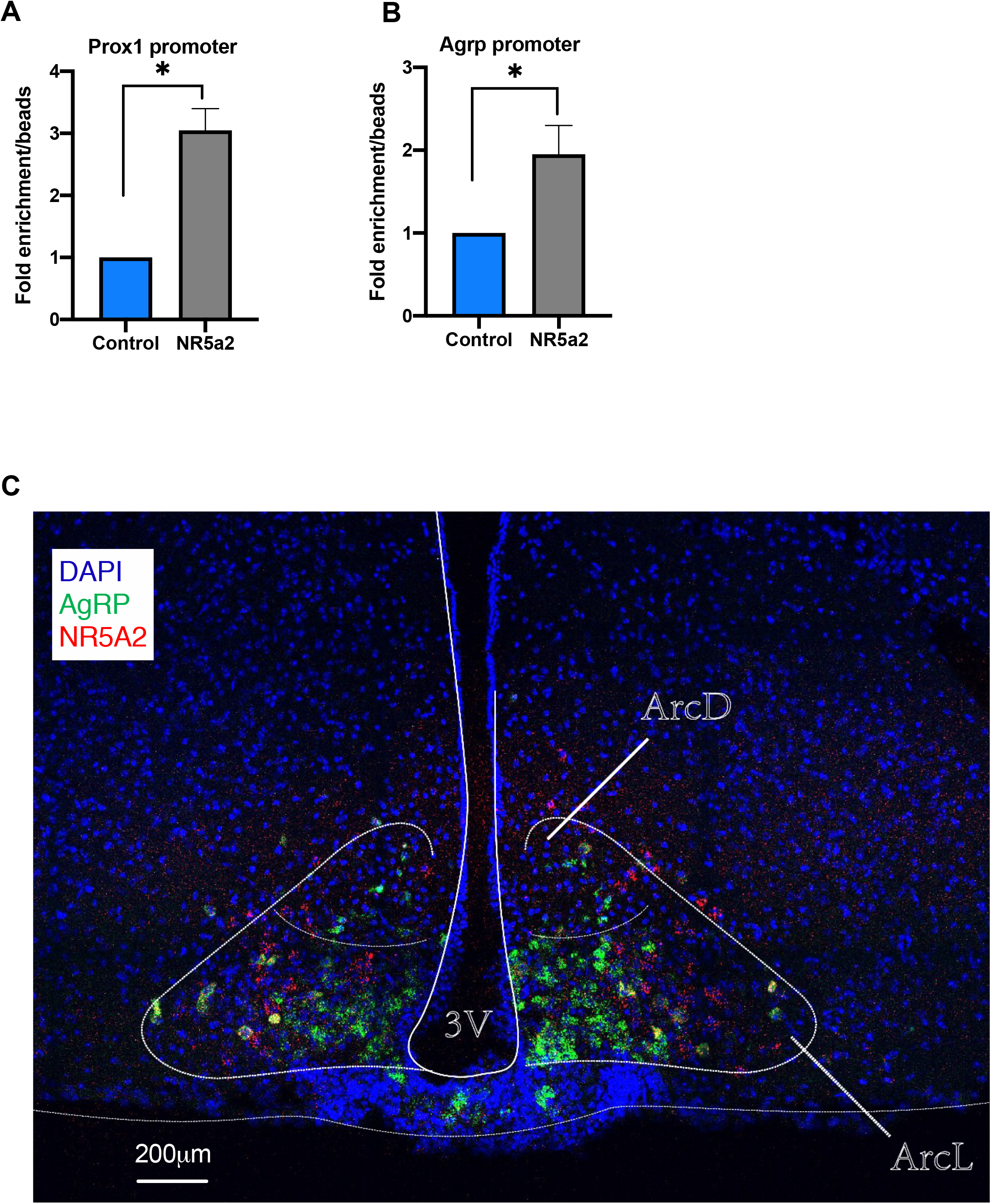
*Agrp* is a direct transcriptional target of Nr5a2. Chromatin immunoprecipitation with Nr5a2 antibodies followed by PCR (chIP-PCR) in the hypothalamic cell line (mHypo-59A) result in enrichment of binding to the (**A**) *Prox1* promoter (positive control) and (**B**) *Agrp* promoter region compared with beads. Data is expressed as mean ± SEM and was analyzed using students t-test, (n= 3 replicates per group), * denotes statistical significance at *p* < 0.05. **C.** In-situ hybridization reveals NR5a2 (red) is specifically localized to the arcuate nucleus (*Agrp*, green) and co-expressed in a subset of *Agrp* neurons (yellow) in the dorsal (D) and Lateral (L) ARC.

## DISCUSSION

In these studies, we used several mouse models to investigate the role of Nr5a2 in AP-induced food intake and weight gain. We first determined that OLZ treatment resulted in a dose-dependent increase in both *Nr5a2* and *Agrp* expression in hypothalamic cells. Furthermore, hypothalamic *Nr5a2* expression was highly induced in mice that were particularly prone to AIWG compared with mice that were relatively protected from AIWG. Administration of SR1848, a specific Nr5a2 inhibitor, decreased OLZ-induced hyperphagia and weight gain and knockdown of *Nr5a2* in the ARC partially reversed OLZ-induced hyperphagia. Importantly, *Agrp* null mice were protected from OLZ-induced hyperphagia and weight gain, despite having elevated hypothalamic *Nr5a2* expression, suggesting this transcription factor may regulate *Agrp* expression. The ChIP-PCR results reported in the current study show, for the first time, that NR5a2 directly binds to the *Agrp* promoter region and suggest that Nr5a2 directly regulates the expression of this pro-feeding neuropeptide in the hypothalamus. Our *in-situ* studies also confirmed that *Nr5a2* and *Agrp* are co-localized in a small subset of cells in the ARC and this co-expression is also further supported by single cell RNA seq studies of the ARC ^65^.

Despite the importance of Agrp in the homeostatic control of feeding, the transcriptional regulation of its expression is still poorly understood. Studies have shown *Agrp* transcription is regulated by key energy sensors, including Peroxisome proliferator-activated receptor gamma coactivator 1-alpha^80^, AMP-activated protein kinase or sirtuin 1, and Estrogen receptor alpha and Signal transducer and activator of transcription 3 ^81^, forkhead box protein O1^82^, Krüppel-like factor 4^83^. Our studies discovered a new transcriptional regulator Nr5a2 to this important list of factors that can regulate Agrp expression. Future chIP-seq studies are warranted to determine the comprehensive transcriptional targets of Nr5a2 in Agrp-expressing neurons. Given that Nr5a2 is also expressed in non-Agrp expressing cells, it will be important to investigate its transcriptional targets in other neuronal population in the ARC.

Nr5a2 has recently been implicated as playing an important role in maintenance of neuronal differentiation and identity in the hippocampus. In these studies, deletion of Nr5A2 in the dentate gryrus cells in vivo lead to a reduction of the number of NeuN as well as Calbindin-positive neurons^84^. Similar studies in the hypothalamus will be necessary to reveal if this a broader function of NR5a2 in mammalian brain function and plasticity.

To enable transcription factors to bind, chromatin must be an ‘open state’ and these accessible regions can be determined using a technique called Assay for Transposase-Accessible Chromatin combined with sequencing (ATAC-seq). ATAC seq studies from the human prefrontal cortex found enriched motifs for NR5a2 target genes in schizophrenia patients (treated with APs) compared with matched case controls ^85^. These studies confirm that APs impact NR5a2 function in the human brain and suggest NR5a2 is an important target for future therapeutic development.

In summary, these studies identify a novel mechanism by which OLZ triggers the transcription of *Agrp* through NR5a2 in a subset of *Agrp* neurons to promote hyperphagia. These findings can be used to inform future clinical development of APs that do not activate hyperphagia and provide deep insights into the regulation of eating behavior. Importantly, it is critical to mitigate AP-induced weight gain to prevent patient non-compliance ^86^ and avoid further exacerbating the growing obesity epidemic and the associated increase in the prevalence of metabolic diseases.

## ACKNOWLEDGMENTS

This study was accomplished through the support by grants including the National Institutes of Health grant R01DK117872 (OO), the National Institute of Health grant P30 DK063491 (Diabetes Research Center, UCSD) (FT and OO) and Larry L. Hillblom Foundation Postdoctoral Fellowship 2019-D-007-FEL (RZ).

## Authors’ contribution

RCZ, ZZ, AP, and OO performed in vivo experiments. RCZ, DZ, PMP, AL performed laboratory analyses. SMC and FT provided intellectual input and oversight of technical expertise. RCZ, OO and FT analyzed the data and wrote the manuscript.

## Conflict of Interest

**None to declare**

## Notes

### Competing Interest Statement

The authors have declared no competing interest.

